# *C. elegans* PVD Neurons: A Platform for Functionally Validating and Characterizing Neuropsychiatric Risk Genes

**DOI:** 10.1101/053900

**Authors:** Cristina Aguirre-Chen, Nuri Kim, Olivia Mendivil Ramos, Melissa Kramer, W. Richard McCombie, Christopher M. Hammell

## Abstract

One of the primary challenges in the field of psychiatric genetics is the lack of an *in vivo* model system in which to functionally validate candidate <u>*n*</u>europsychiatric <u>*r*</u>isk <u>*g*</u>enes (NRGs) in a rapid and cost-effective manner^1^^−^^3^. To overcome this obstacle, we performed a candidate-based RNAi screen in which *C. elegans* orthologs of human NRGs were assayed for dendritic arborization and cell specification defects using *C. elegans* PVD neurons. Of 66 NRGs, identified via exome sequencing of autism (ASD)^4^ or schizophrenia (SCZ)^5^^−^^9^ probands and whose mutations are *de novo* and predicted to result in a complete or partial loss of protein function, the *C. elegans* orthologs of 7 NRGs were found to be required for proper neuronal development and represent a variety of functional classes, including transcriptional regulators and chromatin remodelers, molecular chaperones, and cytoskeleton-related proteins. Notably, the positive hit rate, when selectively assaying *C. elegans* orthologs of ASD and SCZ NRGs, is enriched >14-fold as compared to unbiased RNAi screening^10^. Furthermore, we find that RNAi phenotypes associated with the depletion of NRG orthologs is recapitulated in genetic mutant animals, and, via genetic interaction studies, we show that the NRG ortholog of ANK2, *unc-44*, is required for SAX-7/MNR-1/DMA-1 signaling. Collectively, our studies demonstrate that *C. elegans* PVD neurons are a tractable model in which to discover and dissect the fundamental molecular mechanisms underlying neuropsychiatric disease pathogenesis.

## Introduction

Neuropsychiatric disorders are a group of complex and heterogeneous mental disorders that greatly contribute to human morbidity, mortality, and long-term disability^1^,^11^. Although the genetic architecture of neuropsychiatric disorders, such as schizophrenia, bipolar depression, and autism has been, to varying degrees, refractive to linkage and candidate-gene association studies^11^, recent advances in genomic technologies have paved the way for higher-resolution studies that have identified specific genomic regions or candidate genes that may play key roles in the etiology of these syndromes. Although these approaches, namely common-variant association studies (CVAS)^12^^−^^14^, studies of copy number variation (CNV)^15^, and exome sequencing^4^,^5^,^8^,^16^, have made significant progress toward the identification of DNA sequence variation that may be causal to these disease states, the inability to functionally validate these candidate neuropsychiatric risk genes (NRGs), or model how distinct mutations in these genes impact pathology, has proven prohibitively challenging. This limitation is in striking contrast to other complex diseases, such as cancer or heart disease, where model-organism-based studies have played pivotal roles in gene discovery and validation pipelines, and, ultimately, the development of highly-effective therapeutics. The development of model-organism-based models for studying neuropsychiatric disease, whereby genetic and functional studies could be performed, would enable published and future genomics level data to be leveraged to its full potential.

Because of its inherent simplicity, the *C. elegans* nervous system is an optimal system to perform *in-vivo* functional studies. Its nervous system, which contains approximately 300 neurons, has been fully mapped and is invariant between animals^17^. Notably, anatomical studies of the PVD neuron^10^,^18^^−^^22^, a sensory neuron that responds to harsh mechanical stimuli^23^ and cold temperature^24^, reveal a highly elaborate dendritic arborization pattern that forms a web-like pattern over the majority of the animal. The size and complexity of this neuron has opened the door for detailed studies aimed at revealing the mechanisms underlying the proper development of cell-type-specific dendritic arborization patterns and neuronal development, in general. Accordingly, given that a number of identified NRGs are likely neuronally expressed^3^,^25^, and recent evidence which suggests a role for defective synaptic and/or dendritic morphology in neuropsychiatric disorders^26^,^27^, we used the *C. elegans* PVD neurons as an emerging model in which to carry out *in-vivo* RNAi and genetic studies aimed at revealing the molecular underpinnings of two neuropsychiatric syndromes, autism (ASD) and schizophrenia (SCZ).

In this study, we retrieved 66 NRGs from a literature-based search for ASD-^4^ or SCZ-associated^5^^−^^9^ risk genes. These NRGs were initially identified via the exome sequencing of affected probands, and mutations within these NRGs were classified as *de novo* and predicted to result in a complete or partial loss of protein function. We find that RNAi knockdown of the *C. elegans* orthologs of 7 NRGs resulted in dendritic arborization and/or cell-fate specification defects. We subsequently extended our RNAi studies to include an additional subset of 14 synaptic network-associated (SYN-NET) NRGs, identified via the exome sequencing of SCZ probands and which harbored *de novo* missense mutations that were undefined in terms of impact on protein function^5^.

Collectively, 50 of 80 ASD or SCZ NRGs retrieved from our literature-based search were matched with ≥1 *C. elegans* ortholog, and, in total, the orthologs of 9 of 50 NRGs queried were found to be required for proper PVD cell-fate specification or dendritic arborization. In addition to showing that genetic mutant alleles of a subset of these positive hits phenocopy the observed RNAi-associated defects, we conduct genetic interaction studies which reveal that the cytoskeleton-associated adaptor, *unc-44* (ANK2), is required for SAX-7/MNR-1/DMA-1 signaling, a key signaling pathway central to the proper development of PVD dendritic arbors. In sum, our studies demonstrate the utility of *C. elegans* PVD neurons as a rapid, inexpensive, and powerful model which can be employed to not only functionally validate candidate NRGs, but also serve as a launching point for more extensive genetic studies that may provide key insight into the genetic and molecular mechanisms underlying neuropsychiatric disorders.

## Materials and Methods

### Strains

*C. elegans* strains were maintained on nematode growth media plates at 20°C using standard techniques^28^, unless otherwise noted. The transgenic strain used to visualize PVD neurons in all experiments was NC1687 [*wdIs52 (F49H12.4::GFP) II*]^24^. Additional strains used were: VH624 [*rhIs13 V; nre-1(hd20) lin-15b(hd126) X*]^29^, CB1260 [*unc-44(e1260) IV*], VC1732 [*let-526(gk816) I/hT2 [bli-4(e937) let-?(q782) qIs48] (I:III)*], and the *mnr-1(gof)* strain, EB1649 [*wdIs52 II*; *dzIs43* (*myo-3p::mnr-1a*)]^30^. HML397 [*wdIs52 II; unc-44(e1260) IV*], HML408 [*let-526(gk816) I/hT2 [bli-4(e937) let-?(q782) qIs48] (I:III); wdIs52 II]*, HML429 [*wdIs52 II; rhIs13 V; nre-1(hd20) lin-15b(hd126) X*], and HML478 [*wdIs52 II*; *unc-44(e1260) IV; dzIs43* (*myo-3p::mnr-1a*)] were constructed using standard techniques.

### RNAi screening

RNAi by feeding was performed as described^10^,^29^,^31^ with minor modifications. Briefly, RNAi feeding clones were grown for 15 hours in LB media with 50μg/ml ampicillin. Bacterial cultures were seeded onto agar plates containing 1mM Isopropyl ß-D-1-thiogalactopyranoside (IPTG) and 100μg/ml ampicillin and allowed to dry for 24-36 hours. Three L4-staged wild-type hermaphrodites (strain NC1687) were then placed onto each plate, incubated for 5 days at 20°C, and allowed to lay a brood. After the incubation period, adult progeny were washed off feeding plates with M9 buffer containing 10mM levamisole and mounted onto 2-5% agar pads. For each test, ~20 adult progeny per RNAi clone were scored for the presence of defective PVD dendritic architecture (including a qualitative increase and/or decrease in secondary, tertiary, or quaternary branching or an overt disorganization of the dendritic arbor) or cell specification defects. In cases where the seeding of L4 hermaphrodites onto RNAi plates produced embryonic or larval lethality of the progeny [*pop-1* (*TCF7L2*), *dlg-1* (*DLG1*/*DLG2*), *gpb-1* (*GNB2*), *par-5* (*YWHAZ*)], L1-synchronized animals were placed directly onto RNAi plates, incubated for 3 days at 20°C, and scored at the adult stage. Note that, due to slow growth of the progeny, the incubation period for *daf-21* (*HSP90AA1*), *Y51A2D.7* (*INTS5*), *let-526* (*ARID1B*), *hsp-1* (*HSPA8*), and *unc-10* (*RIMS1*) RNAi was either 5 or 6 days. In addition, RNAi against *daf-21* (*HSP90AA1*), *Y51A2D.7* (*INTS5*), *let-526* (*ARID1B*), *hsp-1* (*HSPA8*), *par-5* (*YWHAZ*), and *lpd-3* (*KIAA1109*) produced a marked decrease in the number of progeny that survived to the adult stage. Therefore, it was not possible to assay ~20 adult progeny per test. For RNAi in the RNAi hypersensitive background (strain HML429), experimental conditions were identical to those conducted using the wild-type strain, except for the minor modification of four L4-staged hermaphrodites being seeded onto RNAi plates. In addition, the incubation period for HML429 animals treated with *ftt-2* (*YWHAZ*) dsRNA was either 5 or 6 days. All RNAi clones in this study were independently tested a minimum of three times, with the exception of *daf-21* and *hsp-1* in the HML429 background. RNAi clones were retrieved from either the Ahringer^32^ or Vidal^33^ RNAi library, and inserts from all positive RNAi clones were sequenced to confirm their identities.

### Plasmid Construction

In addition to clones selected from the Ahringer or Vidal RNAi libraries, RNAi clones *chd-7* (*CHD-8*), *dlg-1* (*DLG1*/*DLG2*), *dpyd-1* (*DPYD*), *nmr-2* (*GRIN2A*), *daf-21* (*HSP90AA1*), and *taf-13* (*TAF13*) were generated by cloning a ~0.7-1.2kb genomic fragment into the RNAi empty vector, pPD129.36. With the exception of the *chd-7* genomic fragment, all genomic fragments were PCR amplified from N2 worm genomic DNA with primers containing the AgeI and KpnI sites at the 5’ and 3’ ends, respectively. For the *chd-7* genomic fragment, primers containing the NotI site at both the 5’ and 3’ ends were used. Primer pairs (without the restriction enzyme sites at the 5’ and 3’ ends) for the *dlg-1*, *nmr-2*, *daf-21*, and *taf-13* RNAi constructs were obtained from the most recent Ahringer RNAi library database that contains the 2012 Supplementary *C. elegans* RNAi plate sets (Source BioScience). Primers used for the construction of the *chd-7* RNAi construct were (5’-3’): ACGCAA<u>GCGGCCGC</u>GAGATGTTTGATAAGGCTTC (forward) and ACGGAA<u>GCGGCCGC</u>ATTTAACATTGAAATCTTCC (reverse). Primers used for the construction of the *dpyd-1* RNAi construct were (5’-3’): AGTTATA<u>CCGGTGCCAGT</u>TGAATAAGTGGGGA (forward, Ahringer primer) and TTTAC<u>AGGTACC</u>TTGGGAAACCATCTAGTTCG (reverse). Restriction enzyme sites are underlined for each of the primers listed. All plasmids were subsequently transformed into the RNase III-deficient *E. coli* strain, HT115(DE3)^34^. Each constructed RNAi plasmid was sequence-verified to confirm its identity.

### Photodocumentation

Defects in PVD dendritic branching were visualized using a Zeiss Axio Scope.A1 microscope equipped with a GFP optical filter set. Images were captured with Spot Advanced Software, Version 5.2, with an added Extended Depth of Focus module.

### Statistics

GraphPad Prism Software, Version 5.0d, was used for all statistical analyses. For each RNAi clone, data from all independent tests were compiled and a weighted percentage (weighted for sample volume) and weighted standard deviation were calculated. RNAi clones were considered positive if the weighted percentage of animals exhibiting PVD arborization defects was statistically different by Fisher’s Exact Test as compared to animals fed the RNAi empty vector, pPD129.36.

### Orthology Assignment

*C. elegans* orthologs of NRGs were identified through the use of the InParanoid^35^,^36^ orthology database (v. 8.0). In cases where InParanoid v. 8.0 did not predict a *C. elegans* ortholog for a specific NRG, InParanoid v. 7.0 predictions, via the OrthoList^37^ database, were used. InParanoid v. 8.0 was used to determine the number of human and *C. elegans* orthologous gene pairs previously tested in an unbiased RNAi screen of *C. elegans* Chromosome IV^10^.

## Results

### *C. elegans* orthologs of human NRGs are enriched for genes required for proper neuronal development

In order to test whether *C. elegans* orthologs of NRGs are required for proper neuronal development, we first selected predicted candidate NRGs via a literature-based search of recent autism^4^ and schizophrenia^5^^−^^9^ exome sequencing studies focused on identifying *de novo* variants. Because RNAi-mediated depletion results in a decrease in transcript levels, thereby leading to a partial to strong loss of protein function, we selected candidate NRGs whose mutations were predicted to be largely gene inactivating. NRGs fitting these criteria were further stratified into a number of variant categories (Supplementary Table 1), namely rLGD, rdNS, RVIS-SCZ, SYN-NET^LGD^, and SYN-NET^MIS^.

Specifically, the <u>*r*</u>ecurrent <u>*L*</u>ikely <u>*G*</u>ene <u>*D*</u>isrupting (rLGD) category (Supplementary Tables 1 and 2) is encompassed by 27 autism-associated^4^ and 2 schizophrenia-associated^5^,^7^ NRGs that harbor nonsense, splice-site, or frameshift mutations in ≥ 2 affected probands. Because so few rLGD targets have been identified in schizophrenia, we sought to also query a number of schizophrenia-associated NRGs that did not explicitly fit into the above, restrictive criteria. One of the additional categories, called <u>*r*</u>ecurrent <u>*d*</u>amaging <u>*N*</u>on-<u>*S*</u>ynonymous (rdNS) (Supplementary Tables 1 and 3), is composed of 21 NRGs^6^,^8^ that 1) harbor a single LGD variant in one proband and a damaging missense variant in ≥ 1 distinct proband(s), or 2) harbor predicted damaging missense variants in ≥ 2 probands. A second additional category includes 16 schizophrenia-associated NRGs^8^ that harbor either a single LGD or broadly damaging missense variant and whose residual variation intolerance score (RVIS) score placed it in the top 15% of genes that, on a population scale, are generally intolerant to mutation (Supplementary Tables 1 and 4). The last two additional variant categories, <u>*Syn*</u>aptic <u>*Net*</u>work-<u>*LGD*</u> (SYN-NET^LGD^) and <u>*Syn*</u>aptic <u>*Net*</u>work-<u>*Mis*</u>sense (SYN-NET^MIS^) (Supplementary Tables 1 and 5), are composed of 19 schizophrenia-associated NRGs that harbor either a single LGD or missense mutation, respectively, and have been predicted to function in a synaptic protein interaction network^5^ using the SynSysNet database. All together, our literature-based search retrieved a total of 27 ASD-associated (Supplementary Table 6) and 40 SCZ-associated (Supplementary Table 7) candidate NRGs whose mutations were predicted (by ≥ 1 algorithm) to eliminate or reduce protein function (Fig. 1a).

**Figure 1.**
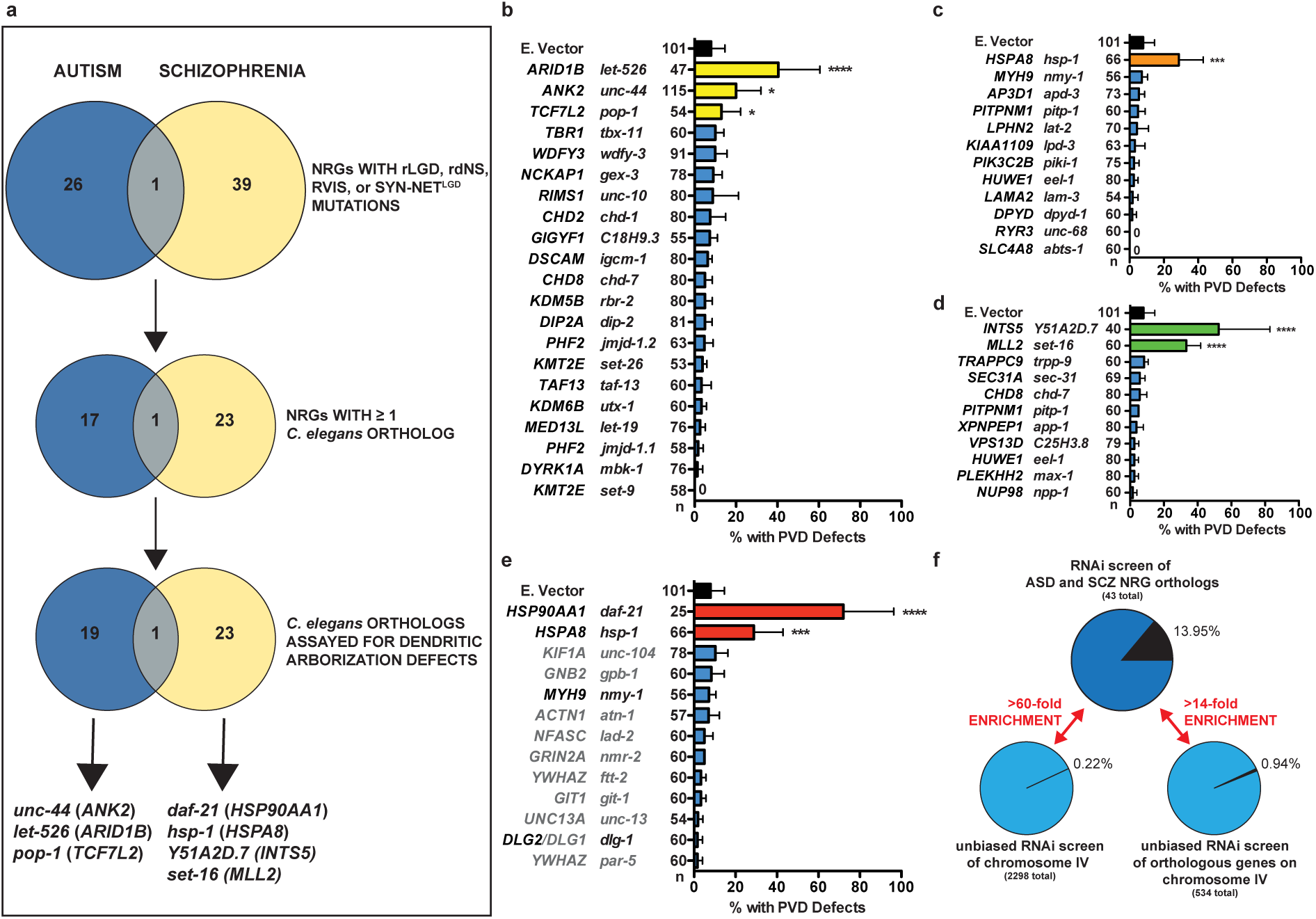
Proper Neuronal Development is Perturbed after the Depletion of *C. elegans* orthologs of ASD or SCZ NRGs. (a) The *C. elegans* orthologs of 66 ASD-or SCZ-associated NRGs were assayed via RNAi, and seven are required for proper neuronal development. A single NRG, CHD8, was present in both the ASD-and SCZ-associated lists. (b-e) RNAi knockdown of the *C. elegans* orthologs of (b) 3/19 rLGD NRGs, (c) 1/12 rdNS NRGs, (d) 2/11 RVIS NRGs, and (e) 2/4 SYN-NET^LGD^ NRGs resulted in PVD dendritic arborization or cell-fate specification defects. Animals exhibiting an overt increase and/or decrease in dendritic branching, a disorganization of the dendritic arbor, or supernumerary PVD cell bodies were scored as defective. For each panel, human NRGs = left column and *C. elegans* orthologs of human NRGs = right column. In panel e, human NRGs, and corresponding *C. elegans* orthologs, labeled in black text represent SYN-NET^LGD^ NRGs, while those labeled in gray text represent SYN-NET^MIS^ NRGs. (f) Selectively assaying *C. elegans* orthologs of ASD and SCZ NRGs results in a >60-fold or >14-fold enrichment as compared to unbiased RNAi screening of 2298 *C. elegans* genes or 534 orthologous gene pairs, respectively. n, number of animals scored. Error bars indicate the weighted standard deviation. \*\*\*\**P*<0.0001, \*\*\**P*<0.001, \**P*<0.05 determined by Fisher’s exact test.

Of the 66 nonoverlapping ASD-and SCZ-associated NRGs harboring predicted deleterious mutations (Fig. 1a, Supplementary Tables 6 and 7), 41 (62%) were identified as having ≥ 1 *C. elegans* ortholog^35^^−^^37^. These 41 NRGs were then matched to a total of 43 *C. elegans* orthologs through the use of the InParanoid orthology database (see methods). In total, the *C. elegans* orthologs of 7 NRGs (Fig. 1a–e and Supplementary Tables 2–5) were found, via RNAi, to be required for two critical aspects of PVD neuronal development: cell fate specification (1 NRG) and dendritic morphogenesis (6 NRGs). We calculated the enrichment of PVD phenotypes associated with our NRG set in two ways. First, as compared to a recently published large-scale RNAi screen of *C. elegans* chromosome IV^10^, in which 2298 *C. elegans* genes were assayed for a role in PVD dendritic morphogenesis, our hit rate of 13.95% (n=6/43) represents a >60-fold enrichment as compared to the hit rate of 0.22% (n=5/2298) associated with unbiased, large-scale RNAi screening (Fig. 1f). As a second metric, we calculated the number of direct human orthologs that are present on chromosome IV and determined that the previously published, unbiased screen assayed a total of 534 orthologous gene pairs. These numbers indicate that the NRG set is enriched >14-fold for phenotypes associated with altered neuronal architecture (Fig. 1f). Notably, although mutations within SYN-NET^MIS^ candidate NRGs are predicted to produce an amino acid change within the encoded protein, these mutations are undefined in terms of whether they are damaging, or broadly damaging, by ≥ 1 algorithm. Therefore, this variant category was not included in the overall hit rate or the enrichment calculation.

### PVD neurons: An assay system to identify NRG-associated perturbations in dendritic patterning and cell fate specification

Based on evidence linking defects in neuronal morphology to neuropsychiatric disorders^26^,^27^,^38^, as well as technical limitations associated with conducting *in vivo* neuronal phenotypic screens in higher-order animal models, we reasoned that the use of the genetically tractable *C. elegans*, the only organism whose complete neuronal wiring diagram has been anatomically mapped^17^,^39^, would be an ideal system in which to elucidate the contribution of NRG orthologs to neuronal development. Specifically, we chose the PVD neurons, the most highly-branched neurons in *C. elegans*, as a triage platform for conducting *in vivo* RNAi screening of NRG orthologs. PVD neurons are a pair of bilaterally symmetric nociceptive neurons that are born at the L2 larval stage^40^. As development proceeds through three larval stages (L2, L3, L4) to the adult stage, primary, secondary, tertiary, and quaternary dendrites are sequentially elaborated at perpendicular angles to one other (Fig. 2a,b,e), giving rise to repeated “menorah” structures that envelop the body of the adult animal^18^^−^^22^ (Fig. 2a). This complex dendritic arborization pattern is in contrast to the simple, unbranched morphology that characterizes the majority of *C. elegans* neurons^17^, and is a unique and advantageous property that allows for the ready identification of dendritic hyperbranching, hypobranching, and self-avoidance phenotypes associated with the loss of NRG ortholog function.

**Figure 2.**
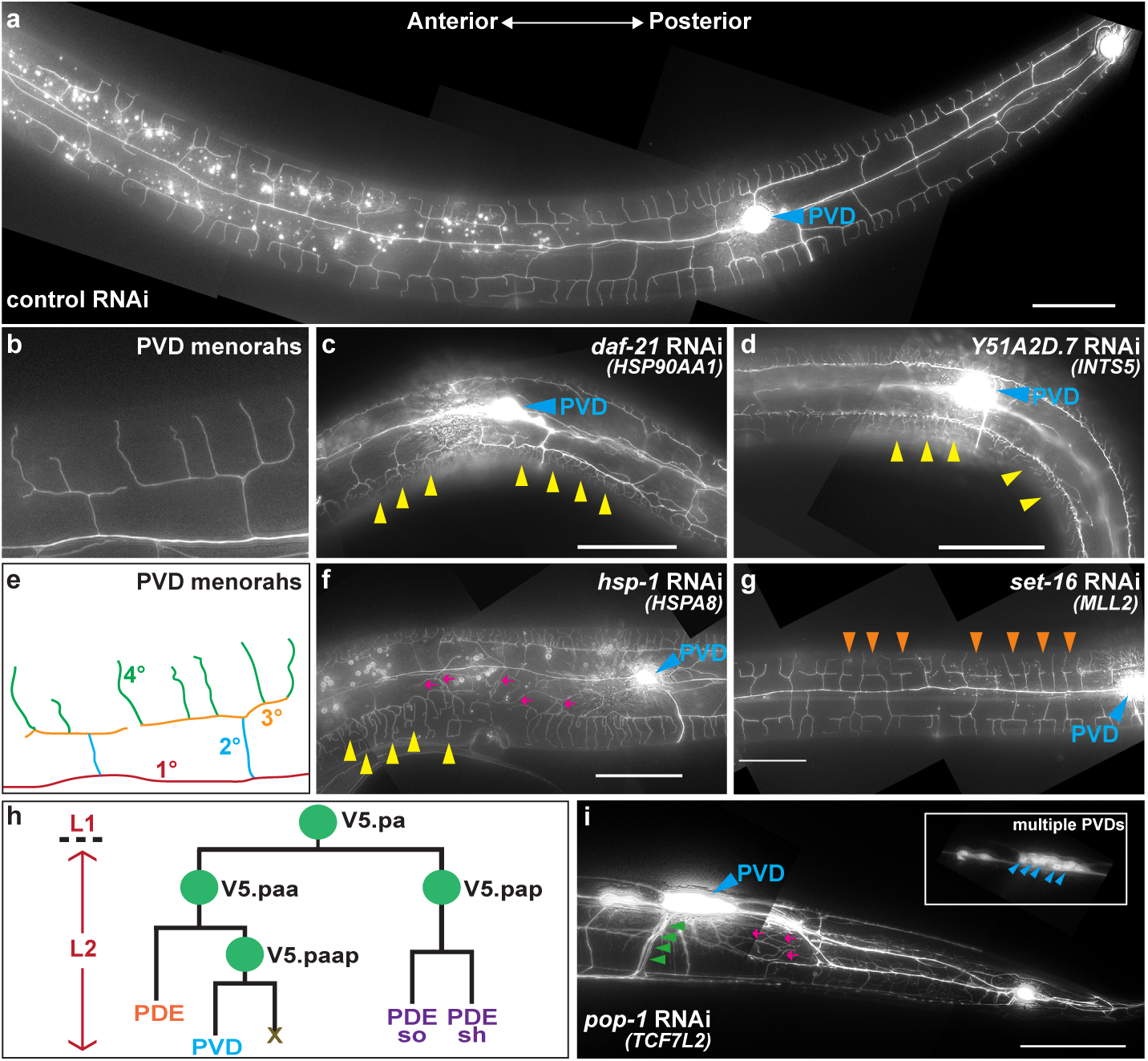
RNAi-mediated Gene Knockdown of NRG Orthologs Disrupts Dendritic Arbor Patterning and Neuronal Cell Fate Specification. (a) PVD neurons form a web-like dendritic arbor that envelops the body of adult-staged animals. (b,e) PVD menorahs are individual dendritic units that sprout ventrally and dorsally from primary dendrites (1°). Each menorah is composed of secondary (2°), tertiary (3°), and quaternary (4°) dendrites. (c,d) RNAi against *daf-21* (*HSP90AA1*) (c) or *Y51A2D.7* (*INTS5*) (d) leads to hyperbranching of 4° dendrites (yellow arrowheads). (f) Animals fed *hsp-1* (*HSPA8*) dsRNA exhibit hyperbranching of 4° dendrites (yellow arrowheads) and increased dendritic branching in the hypodermal region (magenta arrows). (g) RNAi against *set-16* (*MLL2*) results in hypobranching of 4° dendrites (orange arrowheads). (h) cell lineage diagram depicting stage-specific division patterns that give rise to PVD neurons. (i) *pop-1* (*TCF7L2*) dsRNA*-*treated animals exhibit supernumerary PVD cell bodies (blue arrowheads, inset) and axons (green arrowheads) along with an increase in dendritic branching (magenta arrows). In panels c, d, f, g, and i, anteroposterior orientation is as indicated in panel a. In panels a, c, d, f, g, and i, the PVD cell body is labeled with a blue arrowhead. Scale bars: 50μm.

We find that the orthologs of six candidate NRGs play a key role in proper PVD dendritic arbor development. The RNAi knockdown of three NRG orthologs, *daf-21*, *Y51A2D.7*, and *hsp-1*, results in the hyperbranching of secondary, tertiary, and/or quaternary dendrites (Fig. 2c,d,f and Supplementary Tables 3–5). In contrast to a hyperbranching phenotype, animals treated with either *set-16* or *unc-44* dsRNA^10^ exhibit hypobranching of quaternary dendrites and/or reduced quaternary dendrite extension anterior to the PVD cell body (Figure 2g and Supplementary Tables 2 and 4). Interestingly, RNAi depletion of an NRG ortholog may also lead to a mixture of dendritic arborization phenotypes. For example, we find that RNAi knockdown of *let-526* results primarily in a hyperbranching of secondary and tertiary dendrites as well as the misguidance of quaternary dendrites into the hypodermal region, a lateral region that lies between the dorsal and ventral hypodermal/muscle borders (Supplementary Table 2). Other phenotypes that may arise in a minority of *let-526*(RNAi) animals include either dendritic disorganization with defects in self-avoidance or hypobranching of quaternary dendrites (Supplementary Table 2). In addition to dendritic arborization phenotypes, we also identified a single NRG ortholog, *pop-1*, whose RNAi knockdown results in a PVD cell specification defect. In wild-type animals, PVD neurons are derived from the V5 lineage, specifically the V5.pa lineage branch, through a series of repeated cell divisions that take place during the L2 larval stage (Fig. 2h). In *pop-1*(RNAi) animals, defective PVD cell specification leads to supernumerary PVD cells bodies (Fig. 2i). In addition, animals depleted of *pop-1* exhibit dendritic hyperbranching and impaired dendritic self-avoidance (Fig. 2i). Notably, in contrast to the single ventrally-directed PVD axon seen in wild-type animals, *pop-1*(RNAi) animals exhibit multiple ventrally-directed axons (Fig. 2i), suggesting that a partial overlay of PVD dendritic arbors may account for the dendritic hyperbranching and self-avoidance defects. In sum, these findings indicate that NRGs may play functionally distinct roles in controlling developmentally-regulated dendritic arborization and that *C. elegans* PVD neurons can serve as an effective *in vivo* triage platform for screening orthologs of candidate NRGs.

### Depletion of NRG orthologs in an RNAi hypersensitive background reveals additional NRGs that disrupt dendritic arborization

Although PVD neurons in wild-type animals are amenable to RNAi, previous work has shown that the use of RNAi hypersensitive strains to target neuronally-expressed genes enhances RNAi knockdown efficiency^29^,^41^^−^^43^. Accordingly, to complement our RNAi studies in wild-type animals, and potentially identify additional NRG orthologs required for dendritic patterning, we conducted RNAi screening in the *nre-1(hd20) lin15b(hd126)*^*29*^ RNAi hypersensitive background. To conduct these experiments, we focused our screening efforts on the candidate NRGs that comprise the SYN-NET^LGD^ and SYN-NET^MIS^ variant categories.

Through the use of the RNAi hypersensitive strain, we find that the orthologs of two additional candidate NRGs, *YWHAZ* and *GIT1*, are required for proper PVD dendritic arborization (Fig. 3a,d,e and Supplementary Table 8). *YWHAZ* and *GIT1* are represented by three *C. elegans* orthologs, and RNAi knockdown of each results in hyperbranching in the hypodermal region with the phenotypes of *par-5* (*YWHAZ*) (Fig. 3a,d) and *ftt-2* (*YWHAZ*) (Fig. 3a,e) being more penetrant and severe as compared to *git-1* (*GIT1*) (Fig. 3a). In addition, we find that depletion of two candidate NRG orthologs, *daf-21* (*HSP90AA1*) and *hsp-1* (*HSPA8*), leads to a more severe phenotype when knocked down in the *nre-1(hd20) lin15b(hd126)* background as compared to the wild-type strain (Fig. 3a and Supplementary Table 8). Specifically, *daf-21* or *hsp-1* RNAi in the RNAi hypersensitive background results in embyronic lethality (Emb) rather than the dendritic arborization phenotypes observed in wild-type animals, demonstrating the utility of using both wild-type and RNAi hypersensitive strains for NRG ortholog screening. This point is further illustrated when comparing the RNAi hit rate of 15.4% (n=2/13) when using wild-type animals alone (Fig. 3b) versus 38.5% (n=5/13) when using a combination of the wild-type strain and the *nre-1(hd20) lin15b(hd126)* RNAi hypersensitive strain (Fig. 3b). In assessing where SYN-NET^LGD^ and SYN-NET^MIS^-associated positive hits were predicted to function within the postsynaptic terminal^5^ (Fig. 3c), we find that *YWHAZ* acts as a central hub that links the NMDA and AMPA pathways, with *HSP90AA1* and *HSPA8* operating directly upstream and downstream of *YWHAZ*, respectively (Fig. 3c). In addition, *GIT1* is a predicted NMDA pathway component and operates in a distinct, alternate pathway within the postsynaptic terminal (Fig. 3c).

**Figure 3.**
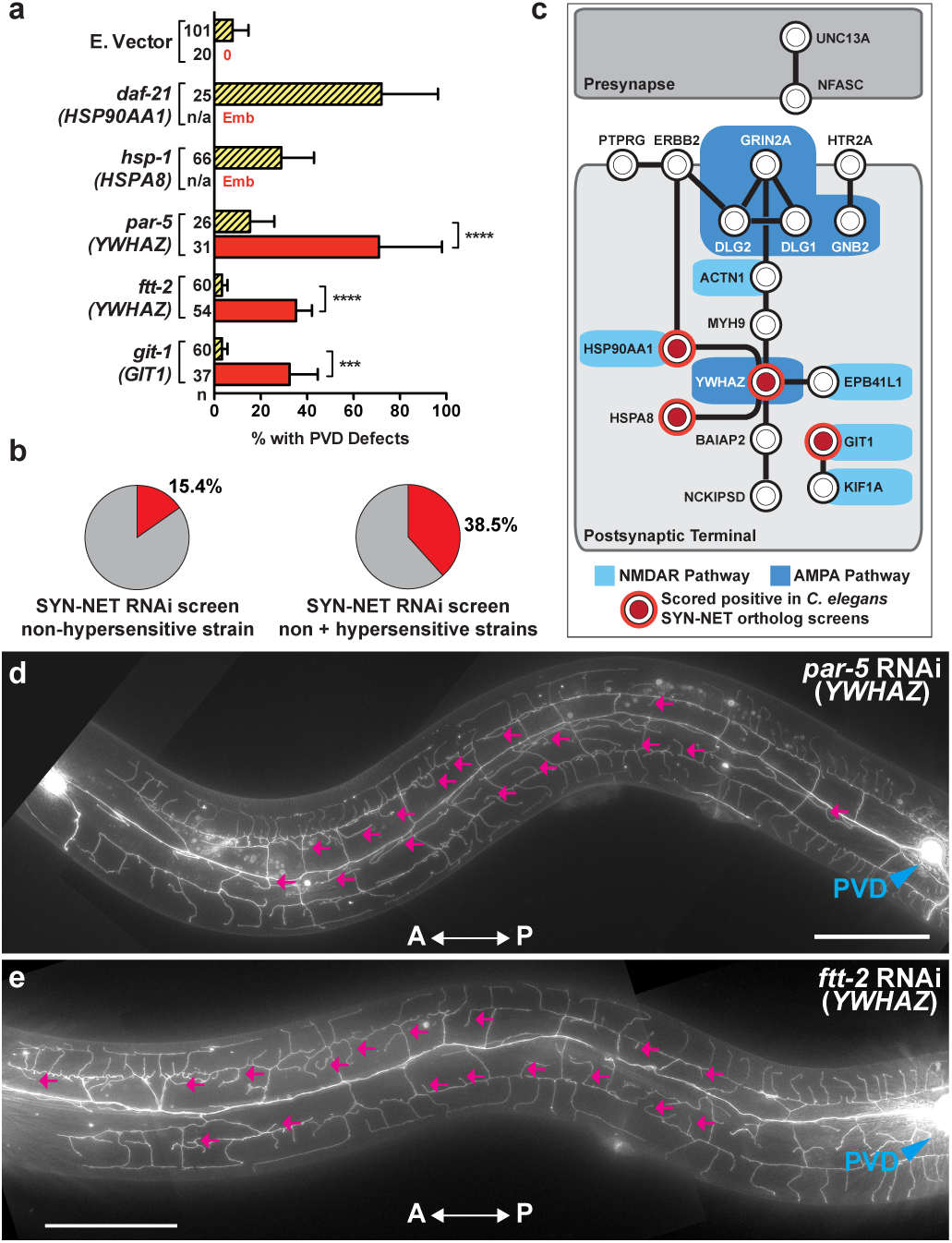
Use of the RNAi Hypersensitive Strain, *nre-1(hd20) lin-15b(hd126)*, offers a Complementary Approach to Identifying Additional NRG Orthologs that Regulate Dendritic Development. (a) RNAi of all SYN-NET NRG (SYN-NET^LGD^ and SYN-NET^MIS^) orthologs identified four NRGs (represented by five *C. elegans* orthologs) as novel regulators of dendritic branching in either the wild-type (yellow bar with diagonal lines) or *nre-1(hd20) lin-15b(hd126)* RNAi hypersensitive (red bar) background. (b) percentage of SYN-NET-associated positive hits identified in the wild-type background (15.4%) as compared to using the wild-type strain in combination with the *nre-1(hd20) lin-15b(hd126)* RNAi hypersensitive strain (38.5%). (c) a synaptic protein-protein interaction network, as reported in Fromer et al.^5^, in which SYN-NET^LGD^ and SYN-NET^MIS^ NRG interactions were identified via the SynSysNet Database (http://bioinformatics.charite.de/synsysnet/). Circles demarcated in red are positive hits identified through the use of the wild-type or the *nre-1(hd20) lin-15b(hd126)* RNAi hypersensitive strain. (d,e) *YWHAZ* orthologs, *par-5* and *ftt-2*, exhibit increased dendritic branching in the hypodermal region (magenta arrows). Anteroposterior orientation is indicated by the white double-headed arrow. In panels d and e, the PVD cell body is labeled with a blue arrowhead. n, number of animals scored. Error bars indicate the weighted standard deviation. \*\*\*\**P*<0.0001, \*\*\**P*<0.001 determined by Fisher’s exact test. Scale bars: 50μm.

### Genetic mutants of NRG orthologs phenocopy the RNAi-induced dendritic arborization defects

To validate dendritic patterning phenotypes associated with the depletion of candidate NRG orthologs, we analyzed genetic mutant alleles of *let-526* and *unc-44*, the *C. elegans* orthologs of *ARID1B* and *ANK2*, respectively. *C. elegans let-526* encodes an ortholog of human *ARID1B*, an ARID (AT-Rich Interacting Domain) domain-containing subunit of the ATP-dependent BAF-B (<u>B</u>RG1/<u>B</u>RM-<u>a</u>ssociated factors; mammalian SWI/SNF) chromatin remodeling complex^44^,^45^ that has been shown to regulate transcriptional output via the repression of hundreds of gene targets^46^. The *let-526*(*gk816*) mutant allele harbors a 1268bp deletion/5bp insertion that disrupts *let-526* isoforms A, C, and D, the same isoforms targeted by the *let-526* RNAi construct. This mutation leads to either the removal of the initiating methionine (isoform A) or to a frameshift (isoforms C and D) and is therefore thought to eliminate their function. As with RNAi-induced depletion of *let-526* (Supplementary Table 2), homozygous *let-526(gk816)* animals exhibit a variety of PVD architectural defects, including hyperbranching, hypobranching, and dendritic arbor disorganization (Fig. 4a,b). Specifically, 22.2% of animals harboring the *gk816* allele exhibit PVD dendritic arbor disorganization (Fig. 4b), 38.9% exhibit dendritic arbor disorganization in combination with hyperbranching and/or hypobranching (Fig. 4b), and 16.7% display dendritic hypobranching alone (Fig. 4b). Interestingly, although *let-526*(*RNAi*) and *gk816* animals exhibit the same classes of dendritic defects, the penetrance of these phenotypes is markedly different, with *let-526(RNAi)* animals primarily exhibiting hyperbranching in the hypodermal region while *let-526(gk816)* mutant animals largely exhibit dendritic arbor disorganization that may be accompanied by hyperbranching and/or hypobranching.

**Figure 4.**
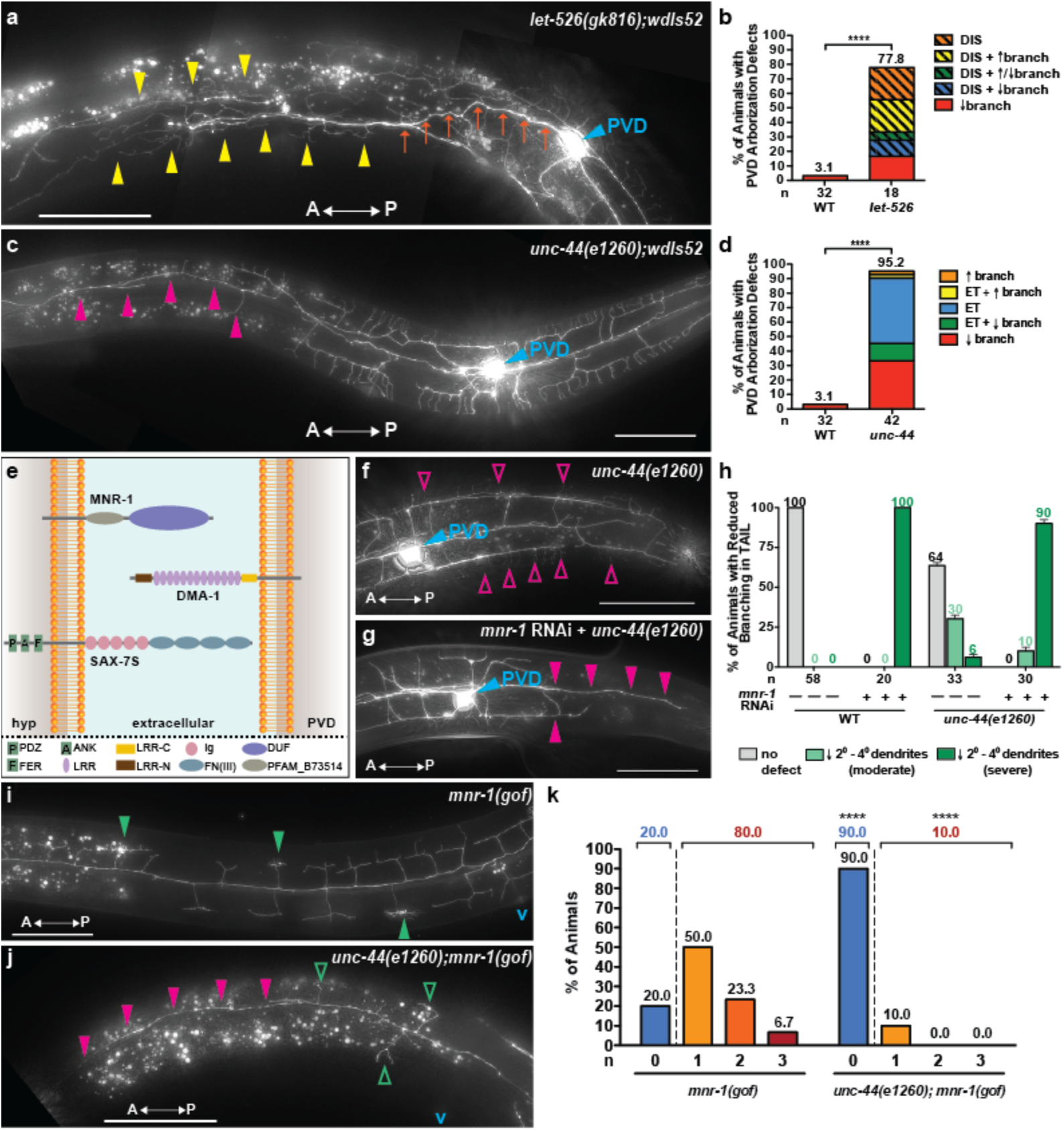
Genetic Mutants Phenocopy RNAi-induced Dendritic Arborization Defects and Genetic Interaction Studies Indicate that *unc-44* (*ANK2*) is Required for SAX-7/MNR-1/DMA-1 Signaling. (a,b) *let-526(gk816)* genetic mutants primarily exhibit a marked disorganization of the dendritic arbor, including spurious branching (yellow arrowheads) and misguidance of the 1° dendrite (orange arrows). A general increase and/or decrease in dendritic branching may also accompany the disorganization phenotype. (b) 22.2% of *let-526(gk816)* mutants exhibit disorganization of the dendritic arbor, while 38.9% exhibit a combination of disorganization and hyper-and/or hypobranching. 16.7% of *let-526(gk816)* mutants exhibit hypobranching alone. (c) In addition to early termination of the 1° anterior dendrite (not shown), *unc-44(e1260)* genetic mutants may also exhibit a loss of 2°, 3°, and 4° dendrites (magenta arrowheads) at the anterior end of the arbor. (d) 33.3% of *unc-44(e1260)* mutants exhibit hypobranching, while 45.2% exhibit early termination of the 1° dendrite. 11.9 % of *unc-44(e1260)* mutants exhibit an early termination of the 1° dendrite in combination with a decrease in dendritic branching. (e) Schematic depicting the SAX-7S/MNR-1/DMA-1 tripartite ligand-receptor complex along with protein domains. (f,h) *unc-44(e1260)* genetic mutants largely retain the ability to form 4° dendrites (magenta open arrowheads) in the tail region (f), although 6% exhibit a complete reduction in 2°, 3°, and 4° dendrites (h). (g,h) A complete reduction in 2°, 3°, and 4° dendrites (magenta arrowheads) in the tail region (g) is exhibited by 90% of *unc-44(e1260)* mutants treated with *mnr-1* dsRNA (h). (i,k) 80% of *mnr-1(gof)* animals elaborate ≥ 1 baobab (green arrowheads) anterior to the vulva, while (j,k) 10% of *unc-44(e1260);mnr-1(gof)* animals sprout ≥ 1 baobab (green open arrowheads) in this region. *mnr-1(gof)* animals harboring the *unc-44(e1260)* allele also exhibit a loss of 2°, 3°, and 4° dendrites (magenta arrowheads) at the anterior end of the PVD arbor. Anteroposterior orientation is indicated by the white double-headed arrow. In panels a, c, f, and g, the PVD cell body is labeled with a blue arrowhead, and “v” in panels i and j marks the approximate location of the vulva. In panels b, d, and h, “n” is the number of animals scored. In panel k, “n” is the number of baobabs present in each genetic background. \*\*\*\**P*<0.0001 determined by Fisher’s exact test. Scale bars: 50μm.

In addition to validating the RNAi phenotype of *let-526*, whose human ortholog, *ARID1B*, has been shown to be a global transcriptional regulator, we also sought to validate RNAi phenotypes associated with orthologs of candidate NRGs that encode cytoskeletal proteins, a class that is directly involved in intracellular organization and transport and one that plays a critical role in cell-cell communication. Among the list of candidate NRGs surveyed in our study, the *C. elegans* ortholog of the cytoskeleton-associated NRG, *ANK2*, was identified in a previous large-scale RNAi study of ours^10^ as being a novel regulator of PVD dendritic arborization. The *C. elegans* ortholog of *ANK2*, *unc-44*, encodes a set of ankyrin-like proteins^47^ that play key roles in axon outgrowth and guidance^48^^−^^50^, neuronal positioning^51^, and axon-dendrite trafficking^52^. Consistent with our previous *unc-44* RNAi findings, we find that animals harboring either the *unc-44(e1260)* or *unc-44(e362)* putative loss-of-function alleles exhibit PVD dendritic hypobranching. In genetic mutant animals, however, the penetrance and severity of the hypobranching phenotype is increased (Fig. 4c,d), and, in addition, an early termination (ET) of the primary dendrite, or a combination of the two phenotypes, is prevalent (Fig. 4d). In contrast to a selective loss of PVD quaternary dendrites in 20% (n=115) of *unc-44(RNAi)* animals^10^ (Fig. 1b and Supplementary Table 2), 33.3% (n=14/42) (Fig. 4d) of *unc-44(e1260)* adult-staged animals exhibit a decrease in PVD dendrite branching anterior to the PVD cell body, with 57% (n=8/14) of this cohort exhibiting a complete loss of secondary, tertiary, and quaternary dendrites in the distal region (anterior to the vulva) of the PVD arbor. We also find that an additional 11.9% (n=5/42) (Fig. 4d) exhibit both dendritic hypobranching distal to the PVD cell body and early termination of the primary anterior and/or posterior dendrite, while 45.2% (n=19/42) (Fig. 4d) exhibit early termination of the primary dendrite alone (n=18/19, ET of primary anterior dendrite; n=1/19, ET of primary anterior and posterior dendrite). Furthermore, we find that a small percentage of animals (4.8%, n=2/42) exhibit either dendritic hyperbranching or a combination of hyperbranching and early termination of the primary anterior dendrite (Fig. 4d). In analyzing genetic mutants harboring the *e362* allele, we observed the same range of PVD dendritic arborization phenotypes as those seen in *unc-44(e1260)* genetic mutants (data not shown). Interestingly, as compared to our previous report, in which *unc-44* RNAi conducted in a wild-type background resulted in hypobranching of quaternary dendrites^10^, we find that animals treated with *unc-44* dsRNA in the *nre-1(hd20) lin15b(hd126)* RNAi hypersensitive background exhibit more penetrant and severe dendritic hypobranching phenotypes (Supplementary Figure 1) that more faithfully recapitulate those observed in *unc-44* genetic mutant animals.

### Genetic interaction studies reveal that *unc-44* (ANK2) is required for SAX-7/MNR-1/DMA-1 signaling

Recent studies have demonstrated that proper PVD dendritic arborization is dependent on the SAX-7/MNR-1/DMA-1 tripartite ligand-receptor complex^30^,^53^,^54^, in which DMA-1 functions as a PVD cell-surface receptor^54^ for the hypodermally-expressed co-ligands, SAX-7S and MNR-1^30^,^53^ (Fig. 4e). Notably, it has been demonstrated that animals harboring a null allele of any component of this ligand-receptor complex exhibit a severe PVD dendritic hypobranching phenotype^30^,^53^,^54^. Because *unc-44* genetic mutant animals also exhibit a reduced-branching phenotype, we attempted to probe whether *unc-44* acts cooperatively with the *sax-7/mnr-1/dma-1* complex by depleting *mnr-1* in an *unc-44(e1260)* genetic mutant background. In contrast to 6% (n=2/33) of *unc-44(e1260)* mutants treated with the L4440 empty RNAi vector (Fig. 4h), 90% (n=27/30) of *mnr-1 (RNAi);unc-44(e1260)* animals exhibit a severe reduction in higher-order (secondary, tertiary, quaternary) branching in the posterior PVD arbor (tail region) (Fig. 4g,h). In addition, the percentage of *mnr-1(RNAi);unc-44(e1260)* animals (10%, n=3/30) exhibiting a moderate decrease in higher-order branching (Fig. 4h) is markedly reduced as compared to *unc-44(e1260)* mutants fed L4440 (30%, n=10/33) (Fig. 4h), further confirming a shift in the severity of the phenotype. Because 100% (n=20) of wild-type animals treated with *mnr-1* dsRNA exhibit a severe decrease in dendritic branching (Fig. 4h) that is identical to that observed in *mnr-1 (RNAi);unc-44(e1260)* animals, these data demonstrate that *mnr-1* is epistatic to *unc-44* and suggest that *mnr-1* and *unc-44* genetically interact to regulate PVD dendritic branch formation.

In order to further test whether *unc-44* is required for *sax-7*/*mnr-1*/*dma-1* signaling, we took advantage of a recently constructed *mnr-1* gain-of-function (gof) strain, in which MNR-1 is ectopically expressed in muscle cells while continuing to be expressed in the hypodermis at endogenous levels^30^. Ectopic expression of MNR-1 in this tissue leads to the formation of “baobabs”, defective PVD menorahs characterized by highly-disorganized and tangled quaternary dendrites that are directly apposed to the MNR-1-expressing muscle cells^30^. We find that *unc-44(e1260*);*mnr-1(gof)* animals exhibit a marked reduction of both wild-type menorahs and baobab structures anterior to the vulva (Fig. 4i,j,k). Quantification of this phenotype reveals that 80% (n=24/30) of *mnr-1(gof)* animals elaborate ≥ 1 baobab anterior to the vulva, while 20% (n=6/30) are devoid of baobab structures (Fig. 4i,k). In contrast, 10% (n=2/20) of *unc-44(e1260);mnr-1(gof)* animals elaborate ≥ 1 baobab anterior to the vulva, while a complete absence of baobabs is observed in 90% (n=18/20) of animals (Fig. 4j,k). These data demonstrate that *unc-44* and *mnr-1* are genetic interacting partners and provides additional evidence in support of the hypothesis that *unc-44* is required for *sax-7/mnr-1/dma-1* signaling. All together, these genetic interaction studies serve to underscore the key roles that cytoskeleton-associated proteins, in conjunction with the *sax-7/mnr-1/dma-1* ligand-receptor signaling complex, play in PVD dendritic arbor development.

## Discussion

It can be easily argued that next generation sequencing technologies will revolutionize progress aimed at understanding complex diseases. While the implementation of genomic sequencing strategies to identify genetic variants associated with ASD and SCZ are timely, a major limitation for harnessing these genomics-level data is the difficulty in modeling ASD or SCZ in model organisms, or *in vitro*, and integrating candidate NRGs into an existing molecular and/or systems-level framework.

Although murine animal models of monogenic forms of ASD, such as Rett Syndrome (RTT) and Fragile X Syndrome (FX) have successfully identified neuronal phenotypes associated with the loss of *Mecp2* (RTT) or *Fmr1* (FX), these monogenic forms of ASD lie squarely in the minority, with ASD and SCZ being largely regarded as not only polygenic, but also multifactorial. Importantly, the sheer number of ASD-or SCZ-associated variants, identified through high-throughput sequencing technologies, makes the issue of prioritizing, validating, and characterizing candidate NRGs via higher-order animal models a major bottleneck. Moreover, there are additional pertinent challenges to utilizing higher-order animal models to study the neurobiological mechanisms of neuropsychiatric disease, including the extraordinary complexity of neuronal connectivity patterns, a lack of neuronal cell type-specific markers that have the ability of track neuronal subtypes throughout animal development, and substantial technical difficulties, particularly in terms of animal viability, in generating single-and multi-gene knockout models aimed at characterizing and ordering genes within genetic pathways.

In recent years, a novel approach for modeling human disease *in vitro*, namely cellular programming and reprogramming technology (CPART), was pioneered^55^,^56^, and recent studies have demonstrated its potential for modeling psychiatric disease. Neuronal cell types generated from the differentiation of human induced pluripotent stem cells^57^^−^^59^ (hiPSC) or from the direct conversion of somatic cells (iNeurons)^60^^−^^62^ can be used to characterize disease-specific neuronal phenotypes (neural differentiation, axon/dendritic/synaptic morphogenesis, electrical and synaptic properties, gene expression) and to study the temporal progression and abnormal development of disease at the cellular level. Notably, when combined with high-throughput genomic sequencing and gene engineering technologies, it can serve as a powerful tool in which to correlate disease-relevant neuronal phenotypes with specific genetic mutations, and holds the promise of revealing key pathophysiological mechanisms and potential therapeutic targets. Important limitations, however, such as a lack of standard operating procedures as it relates to sources of somatic cell types, reprogramming methodology, and metrics to define the pluripotent and differentiated cell states, remain to be resolved. In addition, to more accurately model complex and polygenic diseases, such as ASD or SCZ, three-dimensional (3D) *in vitro* models that recapitulate *in vivo* neuronal morphology and connectivity need to be further developed in order to resolve the contribution of candidate NRGs to cell-autonomous and/or non-cell-autonomous pathways. These roadblocks serve to highlight the continued importance and utility of conducting *in vivo* model-organism based studies to elucidate the genetic, cellular, and molecular mechanisms underlying neuropsychiatric disorders.

Given that the modeling of psychiatric disease in higher-order organisms is extremely challenging, we propose the use of *C. elegans* PVD neurons as an alternate *in vivo* platform for functionally validating and characterizing candidate NRGs. The use of *C. elegans* is ideal for a number of reasons, including 1) its nervous system is simple, containing ~300 neurons, and hard-wired, meaning that neuronal cell position and connectivity are unchanged from animal to animal, 2) due to its transparency, individual neurons and neuronal circuitry can be visualized *in vivo*, and throughout development, through the use of fluorescent transgenic reporters, and 3) it is a powerful genetic model organism in which genetic screens and interaction studies can be used to correlate phenotype to genotype and organize genes within a biological pathway. In addition to *C. elegans* being a rapid, scalable, and inexpensive platform, we reasoned that this combination of attributes, would enable ASD-and SCZ-associated candidate NRGs to be studied in a systematic manner.

As a proof-of-concept, we used candidate-based RNAi screens to query a list of published candidate NRGs identified via exome sequencing and implicated in ASD^4^ or SCZ^5^^−^^9^. We find that RNAi knockdown of the *C. elegans* orthologs of 9 NRGs, spanning all mutational classes defined in this study, resulted in defective PVD dendritic arborization or cell fate specification. Of these 9 NRGs, three are linked to ASD (*ARID1B*, *ANK2*, *TCF7L2*), while six are SCZ-associated (*HSP90AA1*, *HSPA8*, *INTS5*, *MLL2*, *YWHAZ*, *GIT1*). Notably, the variety of phenotypes that were encountered, including dendritic hyperbranching, hypobranching, disorganization, and cell fate specification defects, highlight the utility of PVD neurons as a triage system for assaying the role of candidate NRGs in neuronal development. In addition, we demonstrate that genetic mutants animals harboring mutations in either *let-526* (ARID1B) or *unc-44* (ANK2) phenocopy the observed RNAi defects, and we use genetic interaction studies to show that *unc-44* (ANK2), a cytoskeleton-associated protein, is required for SAX-7/MNR-1/DMA-1 signaling, a key signaling pathway central to the proper development of PVD dendritic arbors.

In sum, our study has focused on the development and implementation of a *C. elegans*-based pipeline (Fig. 5) aimed at determining whether ASD-or SCZ-associated candidate NRG orthologs, identified through large-scale genomics techniques, play a role in the establishment of proper neuronal development. The use of the *C. elegans*-based NRG pipeline to identify and characterize candidate NRG orthologs that are required for various aspects of neuronal morphogenesis can be used as an additional, powerful *in vivo* tool to inform future, more targeted *in vivo* studies in higher-organisms and *in vitro* studies, and, importantly, holds the potential of identifying novel therapeutic targets aimed at ameliorating the effects of neuropsychiatric diseases such as autism and schizophrenia.

**Figure 5.**
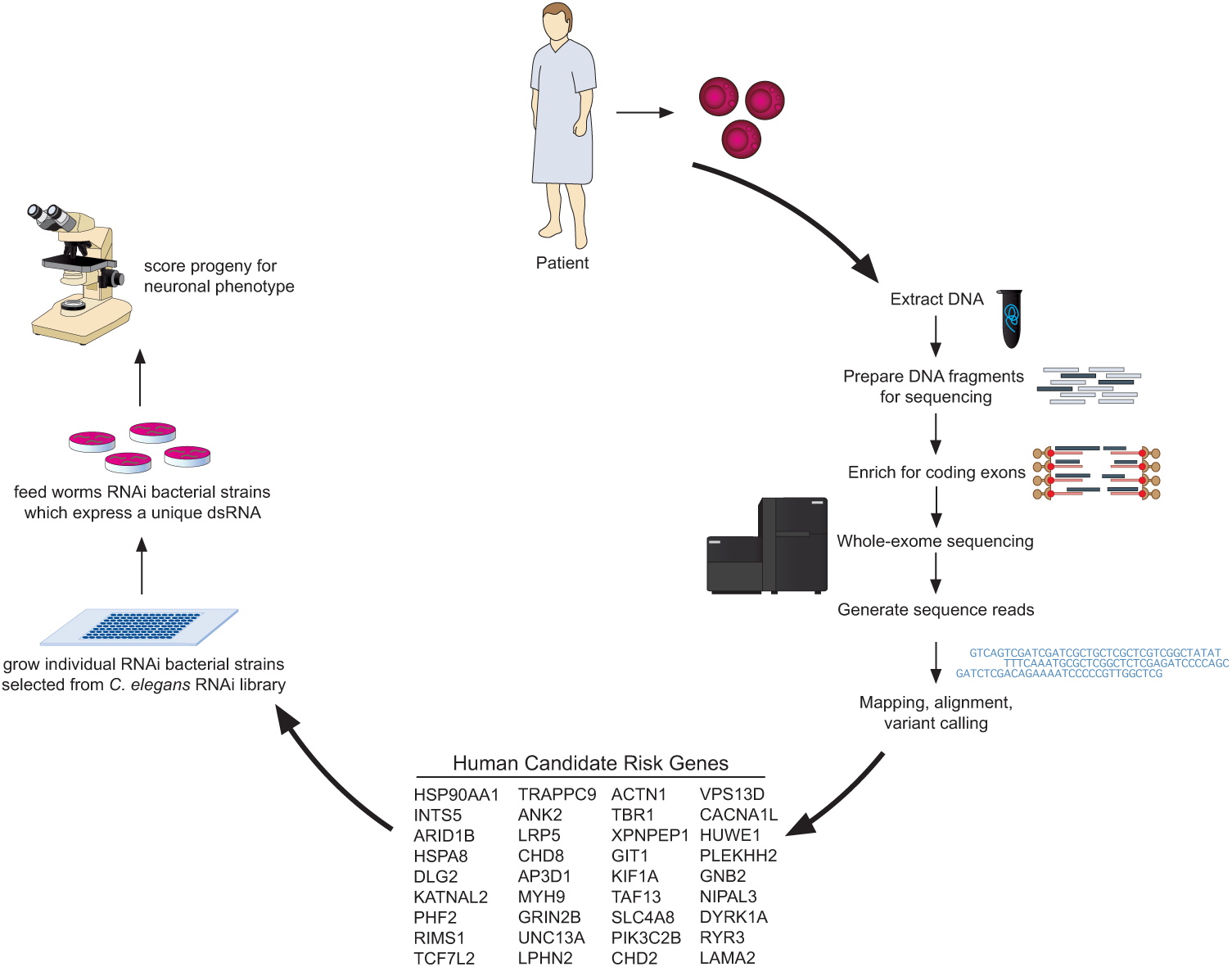
Gene Discovery and Functional Validation Pipeline of Human Neuropsychiatric Risk Genes. After the patient sample is obtained, the process of gene discovery moves forward via rare-variant association studies, such as exome sequencing studies, and the subsequent compilation of human NRGs. Bioinformatic analyses is then used to determine likely *C. elegans* orthologs of the human variants. In this study, we use the *C. elegans* PVD neurons as an *in vivo* triage system to identify and characterize candidate neuropsychiatric risk genes that are required for the proper development of neuronal architecture. These initial validation studies can then be used to guide future, more extensive studies in higher-order model organisms.

## Acknowledgements

We thank the *Caenorhabditis* Genetics Center and Hannes Bülow (Albert Einstein College of Medicine) for *C. elegans* strains; Zaven Kaprielian (Amgen, Albert Einstein College of Medicine), Hannes Bülow (Albert Einstein College of Medicine), and Linda Van Aelst (Cold Spring Harbor Laboratory) for helpful suggestions and critical comments on the manuscript. This work was supported by Cold Spring Harbor Laboratory, the Rita Allen Foundation, and the Simons Foundation Autism Research Initiative grant, SF235988, to Michael Wigler (Cold Spring Harbor Laboratory).

### Conflict of Interest

The authors declare no competing financial interests.

---

**Supplementary Table S1.**
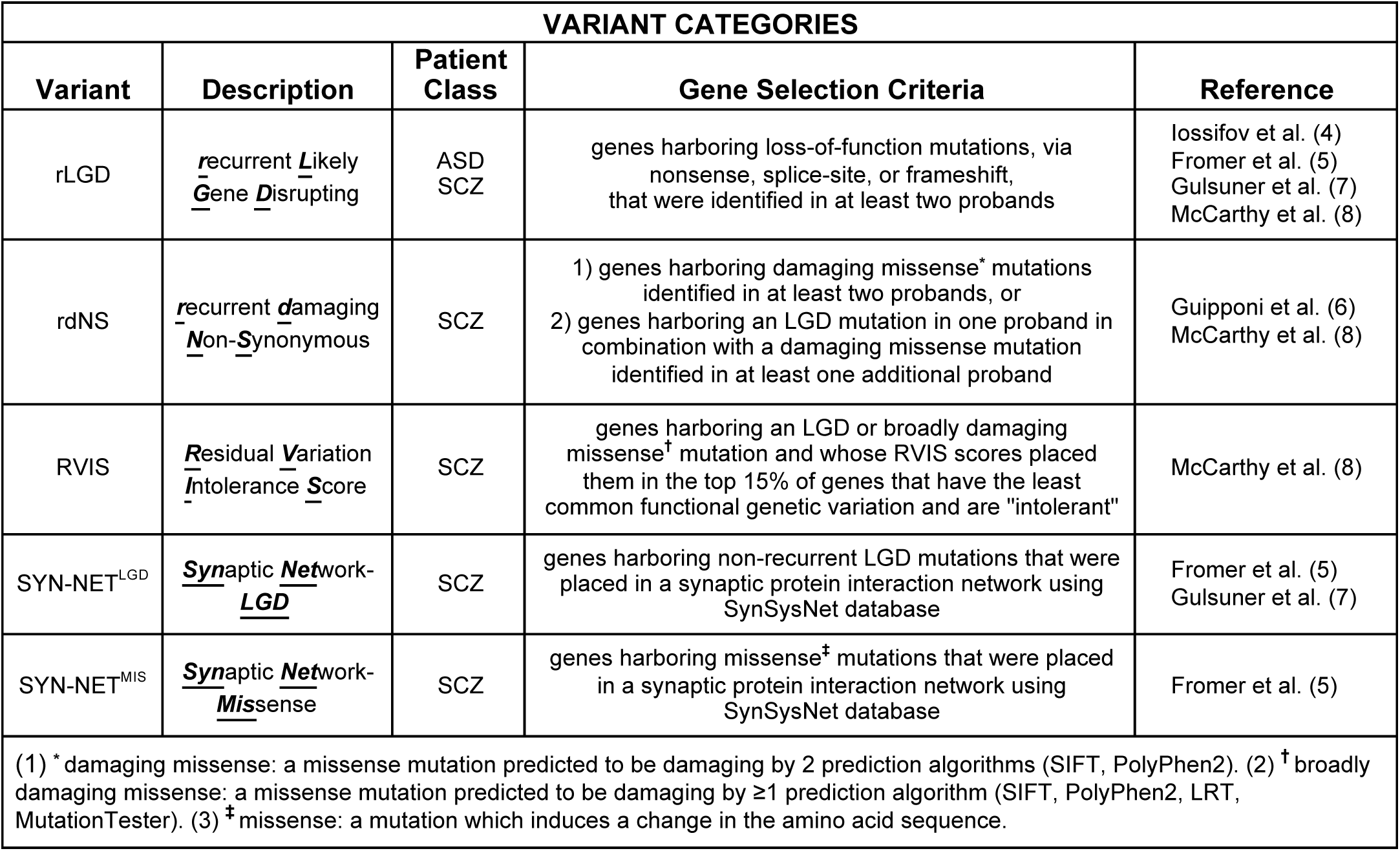
Mutational classes used to categorize ASD-and SCZ-associated NRGs analyzed in this study.

**Supplementary Table S2.** NRGs that lie within the rLGD Mutational Class with corresponding *C. elegans* RNAi data and phenotype descriptions.

| *H. sapiens*<br>Gene | Variant | Ensemble<br>Gene | *C. elegans*<br>Locus ID | *C. elegans*<br>Gene | Animals Defective<br>(% ± SD)<sup>(1)</sup> | n | RNAi<br>Phenotype |
| --- | --- | --- | --- | --- | --- | --- | --- |
| *ARID1B* | rLGD-ASD | ENSG00000049618 | C01G8.9 | *let-526* | 40.4 ± 20.1 | 47 | hyperbranching of dendrites in hypodermal region<sup>(2)</sup><br>hypobanching of quaternary dendrites<br>disorganization of arborization pattern |
| *ANK2*<sup>(3)</sup> | rLGD-ASD | ENSG00000145362 | B0350.2 | *unc-44* | 20.0 ± 12.0 | 115 | hypobanching of quaternary dendrites |
| *TCF7L2* | rLGD-ASD | ENSG00000148737 | W10C8.2 | *pop-1* | 13.0 ± 9.2 | 54 | multiple PVD-like cell bodies |
| *TBR1* | rLGD-ASD | ENSG00000136535 | F40H6.4 | *tbx-11* | 10.0 ± 4.1 | 60 | none |
| *WDFY3* | rLGD-ASD | ENSG00000163625 | C26H9A.2 | *wdfy-3* | 9.9 ± 5.7 | 91 | none |
| *NCKAP1* | rLGD-ASD | ENSG00000061676 | F28D1.10 | *gex-3* | 9.0 ± 4.3 | 78 | none |
| *RIMS1* | rLGD-ASD | ENSG00000079841 | T10A3.1 | *unc-10* | 8.8 ± 12.4 | 80 | none |
| *CHD2* | rLGD-ASD | ENSG00000173575 | H06O01.2 | *chd-1* | 7.5 ± 7.5 | 80 | none |
| *GIGYF1* | rLGD-ASD | ENSG00000146830 | C18H9.3 | *C18H9.3* | 7.3 ± 3.7 | 55 | none |
| *DSCAM* | rLGD-ASD | ENSG00000171587 | F39H12.4 | *igcm-1* | 6.3 ± 2.2 | 80 | none |
| *CHD8* | rLGD-ASD<br>RVIS-SCZ | ENSG00000100888 | T04D1.4 | *chd-7* | 5.0 ± 3.5 | 80 | none |
| *KDM5B* | rLGD-ASD | ENSG00000117139 | ZK593.4 | *rbr-2* | 5.0 ± 3.5 | 80 | none |
| *DIP2A* | rLGD-ASD | ENSG00000160305 | F28B3.4 | *dip-2* | 4.9 ± 3.5 | 81 | none |
| *PHF2* | rLGD-ASD | ENSG00000197724 | F29B9.2 | *jmjd-1.2* | 4.8 ± 4.1 | 63 | none |
| *KMT2E* | rLGD-ASD | ENSG00000005483 | Y51H4A.12 | *set-26* | 3.8 ± 2.2 | 53 | none |
| *TAF13* | rLGD-SCZ | ENSG00000197780 | C14A4.10 | *taf-13* | 3.3 ± 4.7 | 60 | none |
| *KDM6B* | rLGD-ASD | ENSG00000132510 | D2021.1 | *utx-1* | 3.3 ± 2.4 | 60 | none |
| *MED13L* | rLGD-ASD | ENSG00000123066 | K08F8.6 | *let-19* | 2.6 ± 2.5 | 76 | none |
| *PHF2* | rLGD-ASD | ENSG00000197724 | F43G6.6 | *jmjd-1.1* | 1.7 ± 2.4 | 58 | none |
| *DYRK1A* | rLGD-ASD | ENSG00000157540 | T04C10.1 | *mbk-1* | 1.3 ± 2.5 | 76 | none |
| *KMT2E* | rLGD-ASD | ENSG00000005483 | F15E6.1 | *set-9* | 0.0 ± 0.0 | 58 | none |
| *ADNP* | rLGD-ASD | ENSG00000101126 | no ortholog | - |  |  |  |
| *ANKRD11* | rLGD-ASD | ENSG00000167522 | no ortholog | - |  |  |  |
| *FOXP1* | rLGD-ASD | ENSG00000114861 | no ortholog | - |  |  |  |
| *GRIN2B* | rLGD-ASD | ENSG00000273079 | no ortholog | - |  |  |  |
| *KATNAL2* | rLGD-ASD | ENSG00000167216 | no ortholog | - |  |  |  |
| *MKI67* | rLGD-SCZ | ENSG00000148773 | no ortholog | - |  |  |  |
| *POGZ* | rLGD-ASD | ENSG00000143442 | no ortholog | - |  |  |  |
| *SCN2A* | rLGD-ASD | ENSG00000136531 | no ortholog | - |  |  |  |
| *TNRC6B* | rLGD-ASD | ENSG00000100354 | no ortholog | - |  |  |  |
| *WAC* | rLGD-ASD | ENSG00000095787 | no ortholog | - |  |  |  |
| pPD129.36 empty vector | | | | | 7.9 ± 6.7 | 101 | none |

(1) \* = % and standard deviation (SD) denote the weighted % and weighted standard deviation, respectively. (2) <sup>(2)</sup> = "hyperbranching in hypodermal region" may include an increase in secondary and/or ectopic tertiary dendrites as well as a misguidance of quaternary dendrites into the hypodermal region. (3) <sup>(3)</sup> = ANK2 RNAi data as previously reported in Aguirre-Chen et al., 2011.

**Supplementary Table S3.**
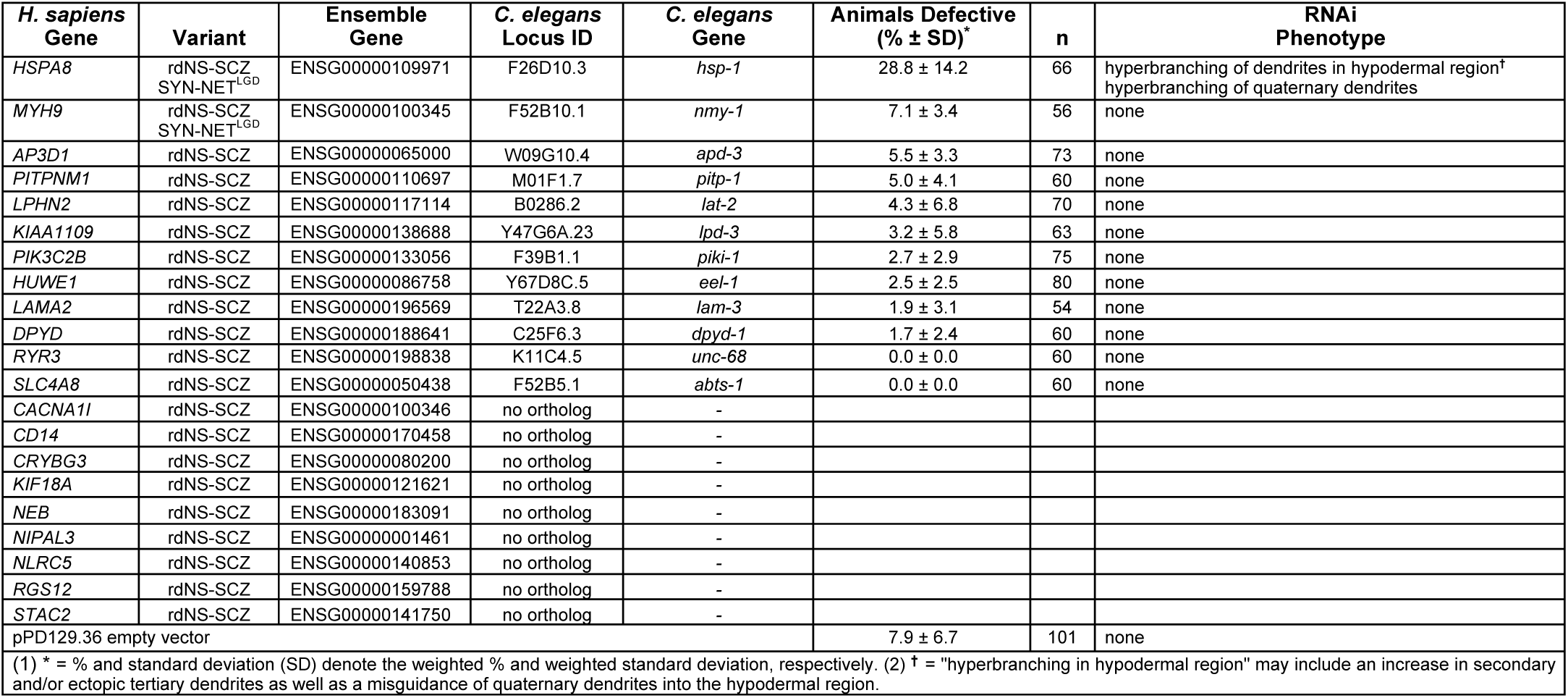
NRGs that lie within the rdNS Mutational Class with corresponding *C. elegans* RNAi data and phenotype descriptions.

**Supplementary Table S4.** NRGs that lie within the RVIS Mutational Class with corresponding *C. elegans* RNAi data and phenotype descriptions.

| *H. sapiens*<br>Gene | Variant | Ensemble<br>Gene | *C. elegans*<br>Locus ID | *C. elegans*<br>Gene | Animals Defective<br>(% ± SD)<sup>*</sup> | n | RNAi<br>Phenotype |
| --- | --- | --- | --- | --- | --- | --- | --- |
| *INTS5* | RVIS-SCZ | ENSG00000185085 | Y51A2D.7 | *Y51A2D.7* | 52.5 ± 30.3 | 40 | hyperbranching of quaternary dendrites |
| *MLL2* | RVIS-SCZ | ENSG00000167548 | T12D8.1 | *set-16* | 33.3 ± 8.5 | 60 | hypobanching of quaternary dendrites<br>reduced quaternary dendrite extension |
| *TRAPPC9* | RVIS-SCZ | ENSG00000167632 | C35C5.6 | *trpp-9* | 8.3 ± 2.4 | 60 | none |
| *SEC31A* | RVIS-SCZ | ENSG00000138674 | T01G1.3 | *sec-31* | 5.8 ± 3.0 | 69 | none |
| *CHD8* | RVIS-SCZ<br>rLGD-ASD | ENSG00000100888 | T04D1.4 | *chd-7* | 5.0 ± 3.5 | 80 | none |
| *PITPNM1* | RVIS-SCZ<br>rdNS-SCZ | ENSG00000110697 | M01F1.7 | *pitp-1* | 5.0 ± 0.0 | 60 | none |
| *XPNPEP1* | RVIS-SCZ | ENSG00000108039 | W03G9.4 | *app-1* | 3.8 ± 4.1 | 80 | none |
| *VPS13D* | RVIS-SCZ | ENSG00000048707 | C25H3.8 | *C25H3.8* | 2.5 ± 2.6 | 79 | none |
| *HUWE1* | RVIS-SCZ<br>rdNS-SCZ | ENSG00000086758 | Y67D8C.5 | *eel-1* | 2.5 ± 2.5 | 80 | none |
| *PLEKHH2* | RVIS-SCZ | ENSG00000152527 | C34B4.1 | *max-1* | 2.5 ± 2.5 | 80 | none |
| *NUP98* | RVIS-SCZ | ENSG00000110713 | ZK328.5 | *npp-10* | 1.7 ± 2.4 | 60 | none |
| *AUTS2* | RVIS-SCZ | ENSG00000158321 | no ortholog | - |  |  |  |
| *LRP5* | RVIS-SCZ | ENSG00000162337 | no ortholog | - |  |  |  |
| *MECP2* | RVIS-SCZ | ENSG00000169057 | no ortholog | - |  |  |  |
| *MKL2* | RVIS-SCZ | ENSG00000186260 | no ortholog | - |  |  |  |
| *RBP3* | RVIS-SCZ | ENSG00000265203 | no ortholog | - |  |  |  |
| pPD129.36 empty vector | | | | | 7.9 ± 6.7 | 101 | none |

(1) \* = % and standard deviation (SD) denote the weighted % and weighted standard deviation, respectively.

**Supplementary Table S5.**
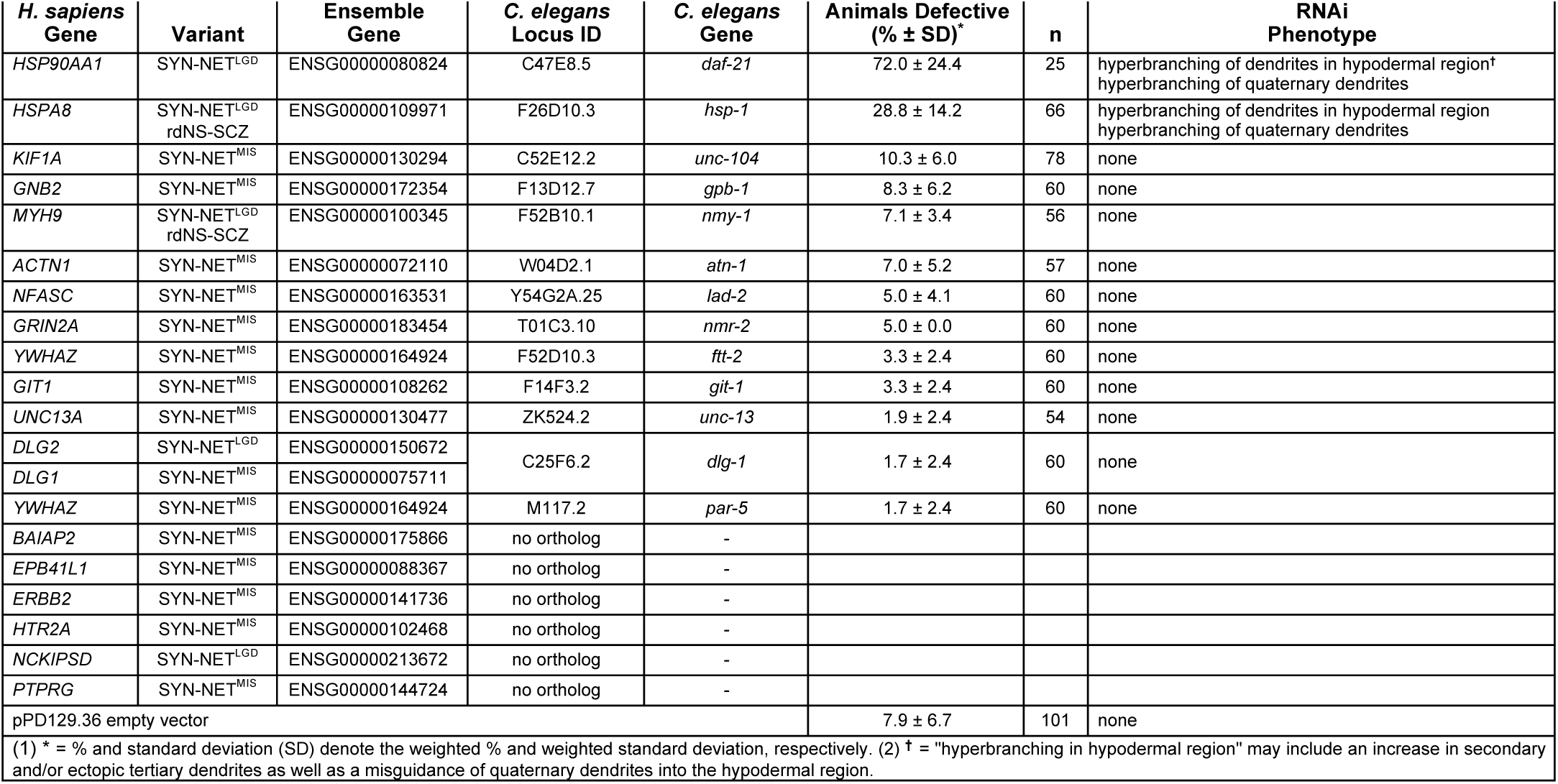
NRGs that lie within the SYN-NET^LGD^ and SYN-NET^MIS^ Mutational Classes with corresponding *C. elegans* RNAi data and phenotype descriptions.

**Supplementary Table S6.** ASD NRGs assayed for defects in neural development.

| AUTISM SPECTRUM DISORDER (ASD):<br>CANDIDATE RISK GENES IDENTIFIED VIA EXOME SEQUENCING |  |  |  |  |
| --- | --- | --- | --- | --- |
| *H. sapiens*<br>Gene | Variant | Ensemble<br>Gene | *C. elegans*<br>Locus ID | *C. elegans*<br>Gene |
| ANK2 | rLGD-ASD | ENSG00000145362 | B0350.2 | *unc-44* |
| ARID1B | rLGD-ASD | ENSG00000049618 | C01G8.9 | *let-526* |
| CHD2 | rLGD-ASD | ENSG00000173575 | H06O01.2 | *chd-1* |
| CHD8 | rLGD-ASD<br>RVIS-SCZ | ENSG00000100888 | T04D1.4 | *chd-7* |
| DIP2A | rLGD-ASD | ENSG00000160305 | F28B3.4 | *dip-2* |
| DSCAM | rLGD-ASD | ENSG00000171587 | F39H12.4 | *igcm-1* |
| DYRK1A | rLGD-ASD | ENSG00000157540 | T04C10.1 | *mbk-1* |
| GIGYF1 | rLGD-ASD | ENSG00000146830 | C18H9.3 | *C18H9.3* |
| KDM5B | rLGD-ASD | ENSG00000117139 | ZK593.4 | *rbr-2* |
| KDM6B | rLGD-ASD | ENSG00000132510 | D2021.1 | *utx-1* |
| KMT2E | rLGD-ASD | ENSG00000005483 | F15E6.1 | *set-9* |
| KMT2E | rLGD-ASD | ENSG00000005483 | Y51H4A.12 | *set-26* |
| MED13L | rLGD-ASD | ENSG00000123066 | K08F8.6 | *let-19* |
| NCKAP1 | rLGD-ASD | ENSG00000061676 | F28D1.10 | *gex-3* |
| PHF2 | rLGD-ASD | ENSG00000197724 | F43G6.6 | *jmjd-1.1* |
| PHF2 | rLGD-ASD | ENSG00000197724 | F29B9.2 | *jmjd-1.2* |
| RIMS1 | rLGD-ASD | ENSG00000079841 | T10A3.1 | *unc-10* |
| TBR1 | rLGD-ASD | ENSG00000136535 | F40H6.4 | *tbx-11* |
| TCF7L2 | rLGD-ASD | ENSG00000148737 | W10C8.2 | *pop-1* |
| WDFY3 | rLGD-ASD | ENSG00000163625 | C26H9A.2 | *wdfy-3* |
| ADNP | rLGD-ASD | ENSG00000101126 | no ortholog | - |
| ANKRD11 | rLGD-ASD | ENSG00000167522 | no ortholog | - |
| FOXP1 | rLGD-ASD | ENSG00000114861 | no ortholog | - |
| GRIN2B | rLGD-ASD | ENSG00000273079 | no ortholog | - |
| KATNAL2 | rLGD-ASD | ENSG00000167216 | no ortholog | - |
| POGZ | rLGD-ASD | ENSG00000143442 | no ortholog | - |
| SCN2A | rLGD-ASD | ENSG00000136531 | no ortholog | - |
| TNRC6B | rLGD-ASD | ENSG00000100354 | no ortholog | - |
| WAC | rLGD-ASD | ENSG00000095787 | no ortholog | - |

**Supplementary Table S7.** SCZ NRGs assayed for defects in neural development.

| **SCHIZOPHRENIA (SCZ):**<br>**CANDIDATE RISK GENES IDENTIFIED VIA EXOME SEQUENCING** |  |  |  |  |
| --- | --- | --- | --- | --- |
| ***H. sapiens***<br>**Gene** | **Variant** | **Ensemble**<br>**Gene** | ***C. elegans***<br>**Locus ID** | ***C. elegans***<br>**Gene** |
| AP3D1 | rdNS-SCZ | ENSG00000065000 | W09G10.4 | *apd-3* |
| CHD8 | RVIS-SCZ<br>rLGD-ASD | ENSG00000100888 | T04D1.4 | *chd-7* |
| DLG2 | SYN-NET<sup>LGD</sup> | ENSG00000150672 | C25F6.2 | *dlg-1* |
| DPYD | rdNS-SCZ | ENSG00000188641 | C25F6.3 | *dpyd-1* |
| HSP90AA1 | SYN-NET<sup>LGD</sup> | ENSG00000080824 | C47E8.5 | *daf-21* |
| HSPA8 | rdNS-SCZ<br>SYN-NET<sup>LGD</sup> | ENSG00000109971 | F26D10.3 | *hsp-1* |
| HUWE1 | rdNS-SCZ<br>RVIS-SCZ | ENSG00000086758 | Y67D8C.5 | *eel-1* |
| INTS5 | RVIS-SCZ | ENSG00000185085 | Y51A2D.7 | *Y51A2D.7* |
| KIAA1109 | rdNS-SCZ | ENSG00000138688 | Y47G6A.23 | *lpd-3* |
| LAMA2 | rdNS-SCZ | ENSG00000196569 | T22A3.8 | *lam-3* |
| LPHN2 | rdNS-SCZ | ENSG00000117114 | B0286.2 | *lat-2* |
| MLL2 | RVIS-SCZ | ENSG00000167548 | T12D8.1 | *set-16* |
| MYH9 | rdNS-SCZ<br>SYN-NET<sup>LGD</sup> | ENSG00000100345 | F52B10.1 | *nmy-1* |
| NUP98 | RVIS-SCZ | ENSG00000110713 | ZK328.5 | *npp-10* |
| PIK3C2B | rdNS-SCZ | ENSG00000133056 | F39B1.1 | *piki-1* |
| PITPNM1 | rdNS-SCZ<br>RVIS-SCZ | ENSG00000110697 | M01F1.7 | *pitp-1* |
| PLEKHH2 | RVIS-SCZ | ENSG00000152527 | C34B4.1 | *max-1* |
| RYR3 | rdNS-SCZ | ENSG00000198838 | K11C4.5 | *unc-68* |
| SEC31A | RVIS-SCZ | ENSG00000138674 | T01G1.3 | *sec-31* |
| SLC4A8 | rdNS-SCZ | ENSG00000050438 | F52B5.1 | *abts-1* |
| TAF13 | rLGD-SCZ | ENSG00000197780 | C14A4.10 | *taf-13* |
| TRAPPC9 | RVIS-SCZ | ENSG00000167632 | C35C5.6 | *trpp-9* |
| VPS13D | RVIS-SCZ | ENSG00000048707 | C25H3.8 | *C25H3.8* |
| XPNPEP1 | RVIS-SCZ | ENSG00000108039 | W03G9.4 | *app-1* |
| AUTS2 | RVIS-SCZ | ENSG00000158321 | no ortholog | - |
| CACNA1I | rdNS-SCZ | ENSG00000100346 | no ortholog | - |
| CD14 | rdNS-SCZ | ENSG00000170458 | no ortholog | - |
| CRYBG3 | rdNS-SCZ | ENSG00000080200 | no ortholog | - |
| KIF18A | rdNS-SCZ | ENSG00000121621 | no ortholog | - |
| LRP5 | RVIS-SCZ | ENSG00000162337 | no ortholog | - |
| MECP2 | RVIS-SCZ | ENSG00000169057 | no ortholog | - |
| MKI67 | rLGD-SCZ | ENSG00000148773 | no ortholog | - |
| MKL2 | RVIS-SCZ | ENSG00000186260 | no ortholog | - |
| NCKIPSD | SYN-NET<sup>LGD</sup> | ENSG00000213672 | no ortholog | - |
| NEB | rdNS-SCZ | ENSG00000183091 | no ortholog | - |
| NIPAL3 | rdNS-SCZ | ENSG00000001461 | no ortholog | - |
| NLRC5 | rdNS-SCZ | ENSG00000140853 | no ortholog | - |
| RBP3 | RVIS-SCZ | ENSG00000265203 | no ortholog | - |
| RGS12 | rdNS-SCZ | ENSG00000159788 | no ortholog | - |
| STAC2 | rdNS-SCZ | ENSG00000141750 | no ortholog | - |

**Supplementary Table S8.** Data associated with the RNAi knockdown of SYN-NET^LGD^ and SYN-NET^MIS^ *C. elegans* orthologs in a wild-type or *nre-1(hd20) lin-15b(hd126)* RNAi Hypersensitive Background.

| *H. sapiens*<br>Gene | Variant | Ensemble<br>Gene | *C. elegans*<br>Locus ID | *C. elegans*<br>Gene | Animals Defective<br>(% ± SD)<sup>*</sup> - NC1687 | n | Animals Defective<br>(% ± SD)<sup>*</sup> - HML429 | n |
| --- | --- | --- | --- | --- | --- | --- | --- | --- |
| *HSP90AA1* | SYN-NET<sup>LGD</sup> | ENSG00000080824 | C47E8.5 | *daf-21* | 72.0 ± 24.4 | 25 | Emb | n/a |
| *HSPA8* | SYN-NET<sup>LGD</sup><br>rdNS-SCZ | ENSG00000109971 | F26D10.3 | *hsp-1* | 28.8 ± 14.2 | 66 | Emb | n/a |
| *YWHAZ* | SYN-NET<sup>MIS</sup> | ENSG00000164924 | M117.2 | *par-5* | 15.4 ± 10.5 | 26 | 71.0 ± 27.2 | 31 |
| *YWHAZ* | SYN-NET<sup>MIS</sup> | ENSG00000164924 | F52D10.3 | *ftt-2* | 3.3 ± 2.4 | 60 | 35.2 ± 6.8 | 54 |
| *GIT1* | SYN-NET<sup>MIS</sup> | ENSG00000108262 | F14F3.2 | *git-1* | 3.3 ± 2.4 | 60 | 32.4 ± 12.1 | 37 |
| pPD129.36 empty vector |  |  |  |  | 7.9 ± 6.7 | 101 | 0.0 ± 0.0 | 20 |
| (1) <sup>*</sup> = % and standard deviation (SD) denote the weighted % and weighted standard deviation, respectively. |  |  |  |  |  |  |  |  |

**Supplementary Figure S1.**
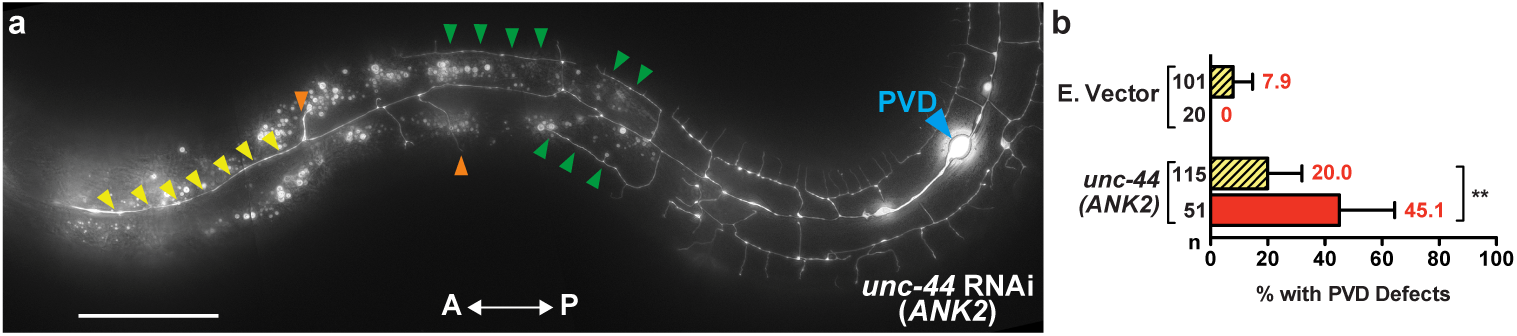
The Penetrance of Dendritic Branching Defects After *unc-44* RNAi is Increased in an *nre-1(hd20) lin-15b(hd126)* RNAi Hypersensitive Background. (a) *unc-44* RNAi knockdown in an *nre-1(hd20) lin-15b(hd126)* background leads to a decrease in 2° (yellow arrowheads), 3° (orange arrowheads), and 4° (green arrowheads) dendrite branch formation. (b) The penetrance of dendritic arborization defects induced by *unc-44* RNAi knockdown increases from 20.0% in a wild-type background^10^ to 45.1% in the *nre-1(hd20) lin-15b(hd126)* RNAi hypersensitive background. Anteroposterior orientation is indicated by the white double-headed arrow, and the PVD cell body is labeled with a blue arrowhead. n, number of animals scored. \*\**P*<0.01 determined by Fisher’s exact test. Scale bar: 50μm.

